# Age differentiation within grey matter, white matter and between memory and white matter in an adult lifespan cohort

**DOI:** 10.1101/148452

**Authors:** Susanne M. M. de Mooij, Richard N. A. Henson, Lourens J. Waldorp, Cam-CAN, Rogier A. Kievit

## Abstract

It is well-established that brain structures and cognitive functions change across the lifespan. A longstanding hypothesis called *age differentiation* additionally posits that the relations between cognitive functions also change with age. To date however, evidence for age-related differentiation is mixed, and no study has examined differentiation of the relationship between brain and cognition. Here we use multi-group Structural Equation Modeling and SEM Trees to study differences *within* and *between* brain and cognition across the adult lifespan (18-88 years) in a large (N>646, closely matched across sexes), population-derived sample of healthy human adults from the Cambridge Centre for Ageing and Neuroscience (www.cam-can.org). After factor analyses of grey-matter volume (from T1- and T2-weighted MRI) and white-matter organisation (fractional anisotropy from Diffusion-weighted MRI), we found evidence for differentiation of grey and white matter, such that the covariance between brain factors decreased with age. However, we found no evidence for age differentiation between fluid intelligence, language and memory, suggesting a relatively stable covariance pattern between cognitive factors. Finally, we observed a specific pattern of age differentiation between brain and cognitive factors, such that a white matter factor, which loaded most strongly on the hippocampal cingulum, became less correlated with memory performance in later life. These patterns are compatible with reorganization of cognitive functions in the face of neural decline, and/or with the emergence of specific subpopulations in old age.

**Significance statement:** The theory of age differentiation posits age-related changes in the relationships between cognitive domains, either weakening (differentiation) or strengthening (de-differentiation), but evidence for this hypothesis is mixed. Using age-varying covariance models in a large cross-sectional adult lifespan sample, we found age-related reductions in the covariance among both brain measures (neural differentiation), but no covariance change between cognitive factors of fluid intelligence, language and memory. We also observed evidence of uncoupling (differentiation) between a white matter factor and cognitive factors in older age, most strongly for memory. Together, our findings support age-related differentiation as a complex, multifaceted pattern that differs for brain and cognition, and discuss several mechanisms that might explain the changing relationship between brain and cognition.

## 1. Introduction

To understand healthy ageing, we must understand the relationship between brain changes and cognitive changes. Although much is known about changes in individual measures such as brain volume or memory performance, less is known about age-related changes in the interrelations *between* neural and cognitive measurements. The *age differentiation* hypothesis describes changes in the organization of cognitive abilities, where differentiation is defined as a low covariance relationship among abilities or factors (Spearman, 1927; Deary & Pagliari, 1990; Hülür, Wilhelm, & Robitzsch, 2011, Blum & Holling, 2017). As people age, there is considerable evidence that they display a loss of differentiation, where cognitive abilities become more correlated, known as *de-differentiation* (Garrett, 1946; Baltes & Lindenberger, 1997; Ghisletta & Lindenberger, 2003; de Frias et al., 2007). However, evidence for this age differentiation-dedifferentiation hypothesis is mixed: Some studies observe a pattern of increase in differentiation followed by de-differentiation (Li et al., 2004), a meta-analysis observed a weak but significant differentiation effect with age (Blum & Holling, 2017), whereas others observe no change in differentiation (Deary et al., 1996; Juan-Espinosa et al., 2002; Zelinski, & Lewis, 2003; Tucker-Drob, 2009; Molenaar et al., 2017). These differences may partly reflect differences in analytical methods, cohorts and sample sizes (Molenaar et al., 2010).

Even less is known about changes in brain organisation as captured by *structural covariance*, meaning the extent to which regional brain structures covary across individuals (Mechelli et al., 2005; Alexander-Bloch, Giedd, & Bullmore, 2013; for brain function, see Park et al., 2004). Previous studies have demonstrated that measures of structural covariance show similarities with structural connectivity and resting-state functional connectivity (Damoiseaux & Greicius, 2009; Seeley et al., 2009; Honey et al., 2009, Alexander-Bloch, Giedd, & Bullmore, 2013; but see Di et al., 2017; Tsang et al., 2017) as well as with developmental trajectories (Zielinski, et al., 2010; Alexander-Bloch et al., 2013). Despite this interest, few studies have used principled methods to investigate whether age-related (de)differentiation occurs for neural measures such as grey matter volume and white matter microstructure. One notable exception is the work by Cox et al. (2016), who found that a single factor for white matter became more prominent with increasing age, suggesting age de-differentiation. A final open question is whether age (de)differentiation occurs not just *within* neural or cognitive domains, but also *between* brain and cognition, such that psychological factors become more or less strongly associated with brain structure across the lifespan.

Understanding the process of age differentiation is crucial for theories of cognitive development and ageing. Older adults may display changes in cognitive strategies: For instance, older individuals may rely more on perceptual salience rather than attentional focus, likely due to poorer internal cues (Lindenberger & Mayr, 2014). Within the neural domain, changes in covariance may reflect a range of important biological processes, including adaptive reorganization (e.g. Cabeza et al., 2002; Greenwood, 2007; Park & Reuter-Lorenz, 2009), regional (Gianaros et al., 2006) or global (Cox et al., 2016) vulnerability to disease states, accumulating structural consequences of lifespan functional connectivity (Seeley et al., 2009), and/or emergence of subgroups that differ in the extent to which they display these patterns.

If age-related changes in cognitive strategy help counter neural decline, then such strategies may eventually induce a more diffuse covariance patterns. For instance, theories of functional plasticity (Greenwood, 2007) and cognitive reserve (Whalley et al., 2004) suggest that adaptive reorganisation in old age leads to decreased covariance between brain structure and cognitive performance. Conversely, theories such as brain maintenance, where preserved cognitive functioning is directly related to maintained brain capacity (e.g. Nyberg et al., 2012), do not predict age-related changes in brain-cognition covariance.

Here we examine age differentiation in a large, healthy, population-derived sample (18-88 years; Cam-CAN, Shafto et al., 2014), using multigroup structural equation modeling (SEM) and SEM-trees. To the best of our knowledge, this is the first study to simultaneously examine age (de)differentiation of grey matter, white matter and cognitive factors.

## 2. Methods

### 2.1 Participants

As part of Phase 2 of the Cambridge Centre for Ageing and Neuroscience (Cam-CAN), data on a wide range of lifestyle, cognitive and neural tests was collected from a healthy, population-based human adult sample, described in more detail in (Shafto et al., 2014). Exclusion criteria include low Mini Mental State Exam (MMSE) (24 or lower), poor hearing (failing to hear 35dB at 1000 Hz in either ear), poor vision (below 20/50 on Snellen test), poor English knowledge (non-native or non-bilingual English speakers), self-reported substance abuse, an indication by the participants’ Primary Care Physician that participation would not be appropriate, and serious health conditions that affect participation (e.g. self-reported major psychiatric conditions, current chemo/radiotherapy, or a history of stroke). We also excluded people with MRI contraindications including disallowed implants, pacemakers, recent surgery or any previous brain surgery, current pregnancy, facial- or very recent tattoos, or a history of multiple seizures or fits) as well as comfort-related contraindications (e.g. claustrophobia or self-reported inability to lie still for an hour). A total of 707 people was recruited for the cognitive assessment (359 females and 348 males) including approximately 100 individuals from each decile (age range 18-88, M=54.63, SD=18.62); usable grey matter was collected from 651 people and white matter from 646 people; sample sizes that are sufficient for moderately complex structural equation models (Wolf et al., 2013). Ethical approval for the study was obtained from the Cambridgeshire 2 (now East of England-Cambridge Central) Research Ethics Committee. Participants gave full informed consent. The raw data to reproduce all analyses can be acquired through the Cam-CAN data portal (https://camcan-archive.mrc-cbu.cam.ac.uk/dataaccess/index.php).

### 2.2 Grey Matter (GM)

To examine grey matter structure, we estimated grey matter volume (GMV) based on the combined segmentation and normalization of 1mm3, T1- and T2-weighted MR images. For more detail on the preprocessing pipeline, see Taylor et al. (2017). We here use GMV for nine ROIs as defined by the Montreal Neurological Institute (Mazziotta et al., 2001). This atlas captures a set of canonical grey matter structures and has a similar number of ROI’s (nine versus ten) as our white matter measure (see below), allowing us to compare evidence for (de)differentiation across grey and white matter using models of comparable complexity. The nine ROIs in the MNI atlas are Caudate, Cerebellum, Frontal Lobe, Insula, Occipital Lobe, Parietal Lobe, Putamen, Temporal Lobe and Thalamus (Figure 1).

**Figure 1.**
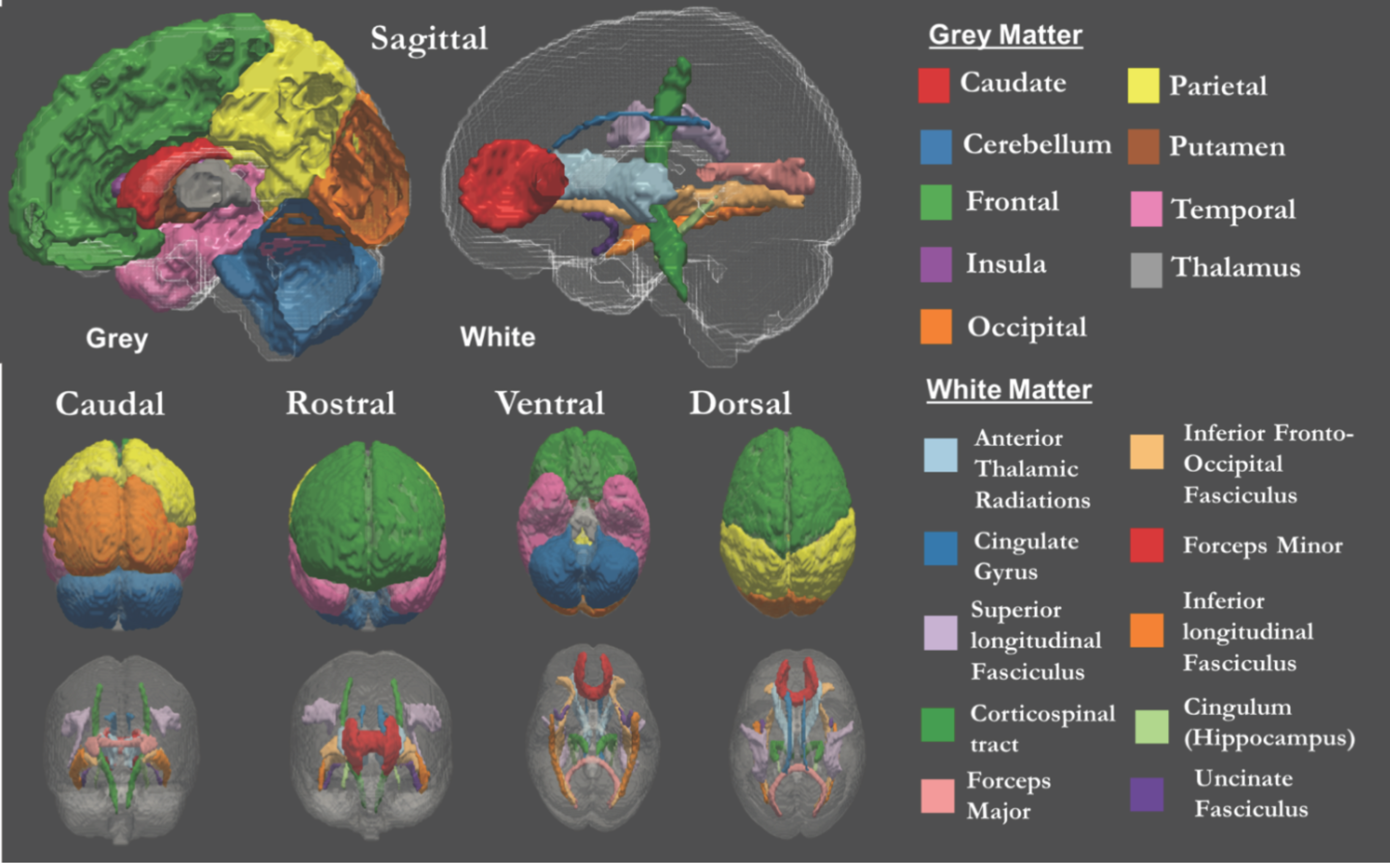
Nine grey and ten white matter tracts as defined by Montreal Neurological Institute (Mazziotta et al., 2001) and Johns Hopkins University white-matter tractography atlas (Hua et al., 2008).

### 2.3 White Matter (WM)

To investigate covariance in white matter structure, we estimated Fractional Anisotropy (FA) values in a set of white-matter ROIs. FA is a measure of the diffusivity of water molecules that is thought to reflect fiber density, axonal diameter and myelination. It is also sensitive to age-related changes in cerebral myelin (Kochunov et al., 2012), although there is discussion on the challenges and limitations of FA (Jones & Cercignani, 2010; Jones, Knösche, & Turner, 2013; Arshad, Stanley, & Raz, 2016; Wandell, 2016). We computed the mean FA for ten ROIs as defined by Johns Hopkins University white-matter tractography atlas (Figure 1; Hua et al., 2008): Anterior Thalamic Radiations (ATR), Cerebrospinal Tract (CST), Dorsal Cingulate Gyrus (CING), Ventral Cingulate Gyrus (CINGHipp), Forceps Major (FMaj), Forceps Minor (FMin), Inferior Fronto-Occipital Fasciculus (IFOF), Inferior Longitudinal Fasciculus (ILF), Superior Longitudinal Fasciculus (SLF) and the Uncinate Fasciculus (UNC). For further details on the white matter pipeline, see Kievit et al. (2016).

### 2.4 Cognitive tasks

Five cognitive tasks were used to assess cognitive processing across three broad cognitive domains: language, memory and fluid intelligence. Language was measured using two tasks: 1) the Spot-the-Word test (Baddeley, Emslie, & Nimmo-Smith, 1993), in which word-nonword pairs (e.g. ‘daffodil-gombie’) are presented and the participant has to decide which is the real word, and 2) a proverb comprehension test, in which participants were asked to provide the meaning of three common proverbs in English (e.g. “Still waters run deep”) yielding a score between 0 and 6. Our measure of fluid intelligence was the standard form of the Cattell Culture Fair, Scale 2 Form A (Cattell, 1971). This pen-and-paper test contains four subsets with different types of abstract reasoning tasks, namely matrices, series completion, classification and conditions. Finally, the third domain memory was assessed using measures of immediate and delayed (after 30-minutes) story recall, as well as recognition, from the logical memory sub-test of the Wechsler Memory Scale Third UK edition (Wechsler, 1997).

### 2.5 SEM Analyses

To improve convergence, prior to the SEM analyses, the neural and cognitive measures were scaled to a standard normal distribution. We used full information maximum likelihood estimation (FIML) and Robust Maximum Likelihood estimator with a Yuan-Bentler scaled test statistic (MLR) to account for violations of multivariate normality. To ensure possible outliers did not affect the results, we fit the models with both full data as well as data treating univariate outliers (z-scores greater than 4 or -4) as missing. Doing so did not affect any model comparison meaningfully, so we report the results for the full dataset. We used SEM to test for evidence for neural and cognitive age (de)differentiation in the following three steps: 1) establish an appropriate measurement model, 2) examine adult lifespan patterns of the factor scores, and 3) formally test for age (de)differentiation using Multigroup confirmatory factor analysis (MGCFA) and SEM trees (see below for more detail).

All models were fit using the package Lavaan (Rosseel, 2012) in the statistical software R (R Core Team, 2016). We assessed overall model fit using the *χ^2^* test, RMSEA and its associated confidence interval, CFI and SRMR (Schermelleh-Engel, Moosbrugger, & Müller, 2003). We considered good fits as follows: RMSEA < 0.05 (0.05-0.08 is acceptable), CFI > 0.97 (0.95-0.97 is acceptable) and SRMR < 0.05 (0.05-0.10 is acceptable). For the MGCFA, we compared models directly with the likelihood ratio test, the AIC and Akaike Weights (Wagenmakers & Farrell, 2004) and the sample size adjusted BIC (saBIC, with associated Schwarz weights). For all age comparisons, we defined three discrete, equally sized subgroups: Young, Middle and Old (see Table 1). For each lifespan multigroup comparison, we compared a model where factor covariance was equality-constrained across the three age groups to a model where they were freely estimated. In the constrained model, all parameters were constrained between the groups, except for the means of the factors (to allow for age-related declines). By comparing these nested models, we could determine whether there is evidence for changing factor covariance structure across the lifespan.

**Table 1:** Demographics of age groups (young, middle and old) for neural and cognitive measures

|  | Age group | Sample size (*N*) | Mean age (*M*) | Standard deviation age (*SD*) |
| --- | --- | --- | --- | --- |
| Grey Matter | Young | 217 | 32.82 | 6.92 |
|  | Middle | 217 | 54.56 | 6.25 |
|  | Old | 217 | 75.56 | 5.91 |
| White Matter | Young | 215 | 32.87 | 6.89 |
|  | Middle | 216 | 54.69 | 6.32 |
|  | Old | 215 | 75.66 | 5.85 |
| Cognition | Young | 235 | 33.10 | 7.19 |
|  | Middle | 236 | 54.86 | 6.37 |
|  | Old | 236 | 75.84 | 5.87 |

In cases where the likelihood ratio test yielded evidence for age differentiation, we visualised the differences by using a technique inspired by Local Structural Equation Models (LSEMs; Hildebrandt, Wilhelm, & Robitzsch, 2009; see also Hülür et al., 2011). This technique allows us to visualize age gradients in model parameters of the covariance structure in a more continuous matter, rather than creating age groups. To do so, we estimated the covariance between factors using a series of age-weighted SEMs for the CFA models with subsets of the sample (N = 260 for WM; N=300 for GM, due to estimation variability) in one-year steps from 18 to 88 years. Next, a kernel function was used to weigh and smooth the observations according to the age gradients (Hildebrandt et al., 2009). The following bandwidth (bw) of the kernel function was used to smooth the age-weighted samples:

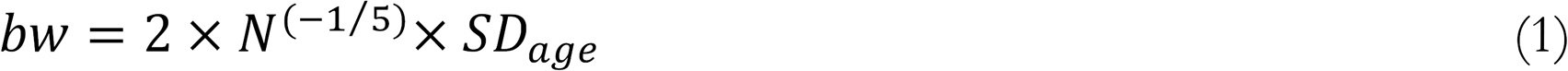

Visualizing factor covariance allowed for the identification of lifespan patterns including differentiation and de-differentiation. If the data are in line with age differentiation, we expect to find that the nested multigroup model with the freely estimated covariance structure is preferred, in such a way that the older subgroup has lower covariance between factors. Evidence for age dedifferentiation would suggest a preference for the freely estimated model, but with higher covariance between the factors in the older subgroup. We first examine differentiation within each domain (grey matter, white matter and cognition), and finally examine brain-cognition covariance differences. Finally, we used Structural Equation Model Trees (SEM Trees), which combine the strength of Structural Equation models (SEM) and decision trees (Brandmaier et al., 2013, 2016). SEM trees partition a dataset repeatedly into subsets based on some covariate(s) of interest to examine whether a likelihood ratio test suggests sufficient evidence of significantly different parameter estimates in each possible subgroup. This method allows us to find covariates and covariate interactions that predict differences in model parameters (in observed and latent space) in a hierarchical fashion. The addition of SEM Trees to the multigroup analyses enables us to analyse age in a continuous nature and trace potential age differences in optimal splits. In this study, SEM Trees were used to investigate whether the covariance structure in the same neural and cognitive factors model as used in the multigroup SEM models changed with age. According to the (de)differentiation hypothesis, SEM Trees would split the dataset into subsets with different covariance structures according to the continuous covariate age.

All SEM Trees were analyzed with the package ‘SEM Trees’ (Brandmaier et al., 2013) in R using on the OpenMx package for SEM. We imposed the same models as with the multigroup SEM to compare the results in favor of or against the differentiation hypothesis. All paths were constrained, except for the covariance between the factors and the factor means to allow age-related decline, but since the factor means change alongside the covariance, the source of the potential split is rather ambiguous. Notably, this technique allows for the specification of focal parameters, such that only differences in model fit due to these key parameters are used to partition the data into subsets. Here, we only base possible splits on the factor covariance, as these reflect the age differentiation hypothesis. The criterion for best split is based on a Bonferroni-corrected likelihood ratio test of differences between the groups resulting from a given split (Brandmaier et al., 2013). To ensure a sufficient number of participants given model complexity, we only allowed splits where the minimal sample per subgroup would be at least 200 participants.

## 3. Results

### 3.1 Grey and white matter covariance

In order to specify a measurement model amenable to multigroup Confirmatory Factor Analysis, we first examined a plausible candidate model using an Exploratory Factor Analysis (EFA). For grey matter, we established a three-factor solution was preferred. This three-factor model showed adequate fit in the following CFA analysis: *χ^2^* (19) = 82.384, p < .001, RMSEA = .072 [.057 .087], CFI = .990, SRMR = .016. For white matter, a three-factor model showed marginally acceptable fit: *χ^2^* (26) = 133.897, p < .001, RMSEA = .080 [.068 .093], CFI = .966, SRMR = .025. The measurement models are shown in Figure 2, along with their correlation matrices. As the precise factors will depend to some extent on the atlas used we will not label the factors, but examine the covariance patterns in more detail. The first grey matter factor (teal) is characterized by strong loadings especially on the insula. The second grey matter factor (blue), is characterized by a relatively broad set of medium sized factor loadings, with an especially strong factor loading for temporal and thalamic grey matter volume. The third grey matter factor (pink) is characterized most strongly by parieto-frontal covariance. Although a single factor model fits poorly for grey matter, the correlations between grey matter factors are relatively strong, especially in comparison to the white matter factors, which are more globally differentiated. The white matter measurement model also yields three factors. The first white matter factor (red) is characterized by strong loadings on more posterior ILF and forceps major tracts, and a negative factor loading on the cingulum. The second white matter factor (yellow) is characterized most strongly by the cingulum, but has a broad set of positive factor loadings across the majority of tracts. Finally, the third factor (green) loads most strongly on the ventral cingulum. The effects of age on the factor scores are shown in Figure 3, revealing different effect sizes, as well as different functional forms (linear and non-linear)

**Figure 2.**
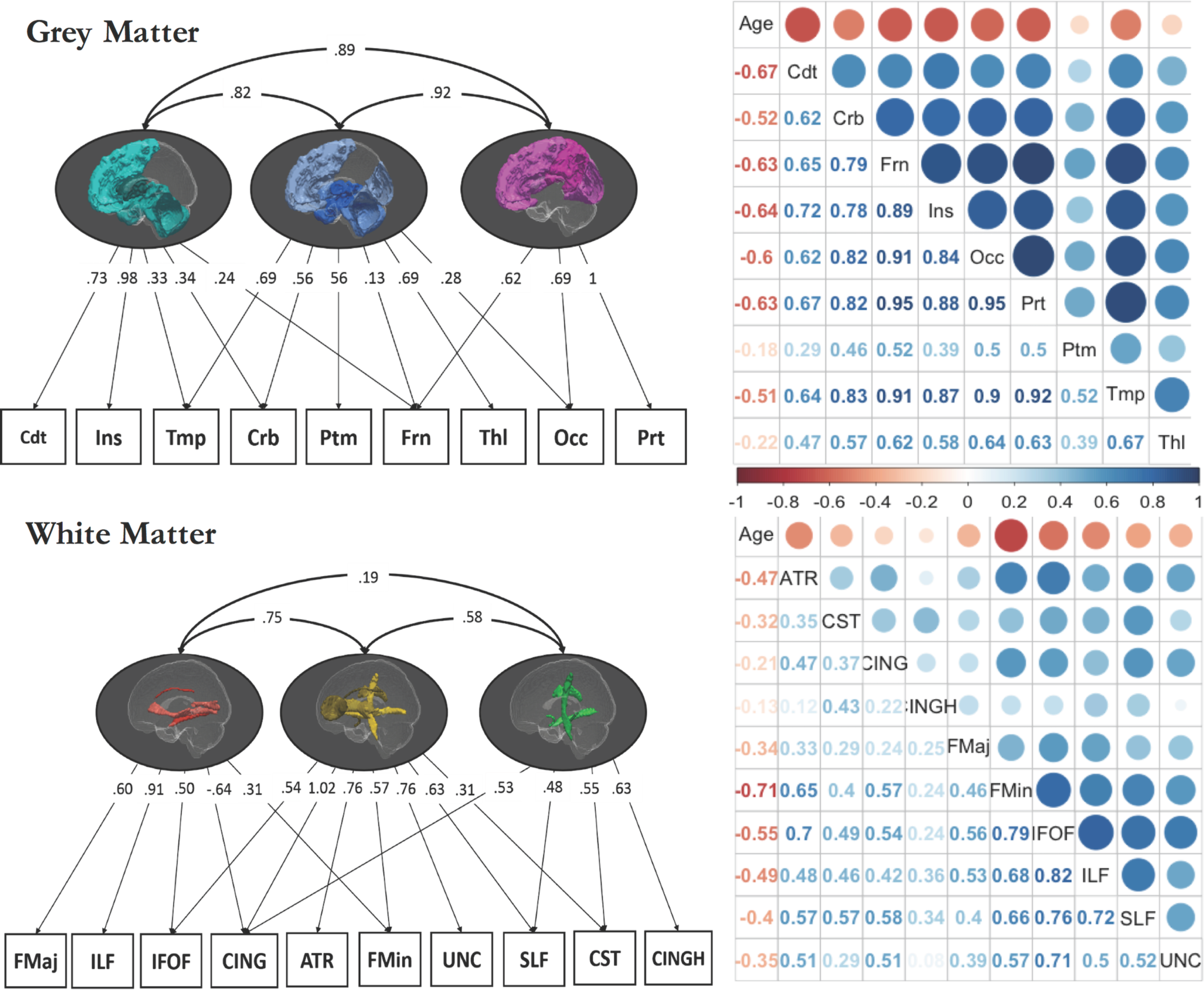
The three-factor model for grey-matter (top left) underlies nine ROIs: Caudate (Cdt), Insula (Ins), Temporal (Tmp), Cerebellum (Crb), Putamen (Ptm), Frontal (Frn), Thalamus (Thl), Occipital (Occ), Parietal (Prt). The three-factor model for white-matter (bottom left) with ten ROIs: Forceps Major (FMaj), Cingulate Gyrus (CING), Inferior Fronto-Occipital Fasciculus (IFOF), Inferior Longitudinal Fasciculus (ILF), Anterior Thalamic Radiations (ATR), Forceps Minor (FMin), Uncinate Fasciculus (UNC), Superior Longitudinal Fasciculus (SLF), Corticospinal Tract (CST), Hippocampal Cingulum (CINGH). The darker colours in the lateral brain views represent the regions with the highest factor loadings. Path coefficients are fully standardized. The correlation matrices are shown for grey matter (top right) and white matter (bottom right), along with age.

**Figure 3.**
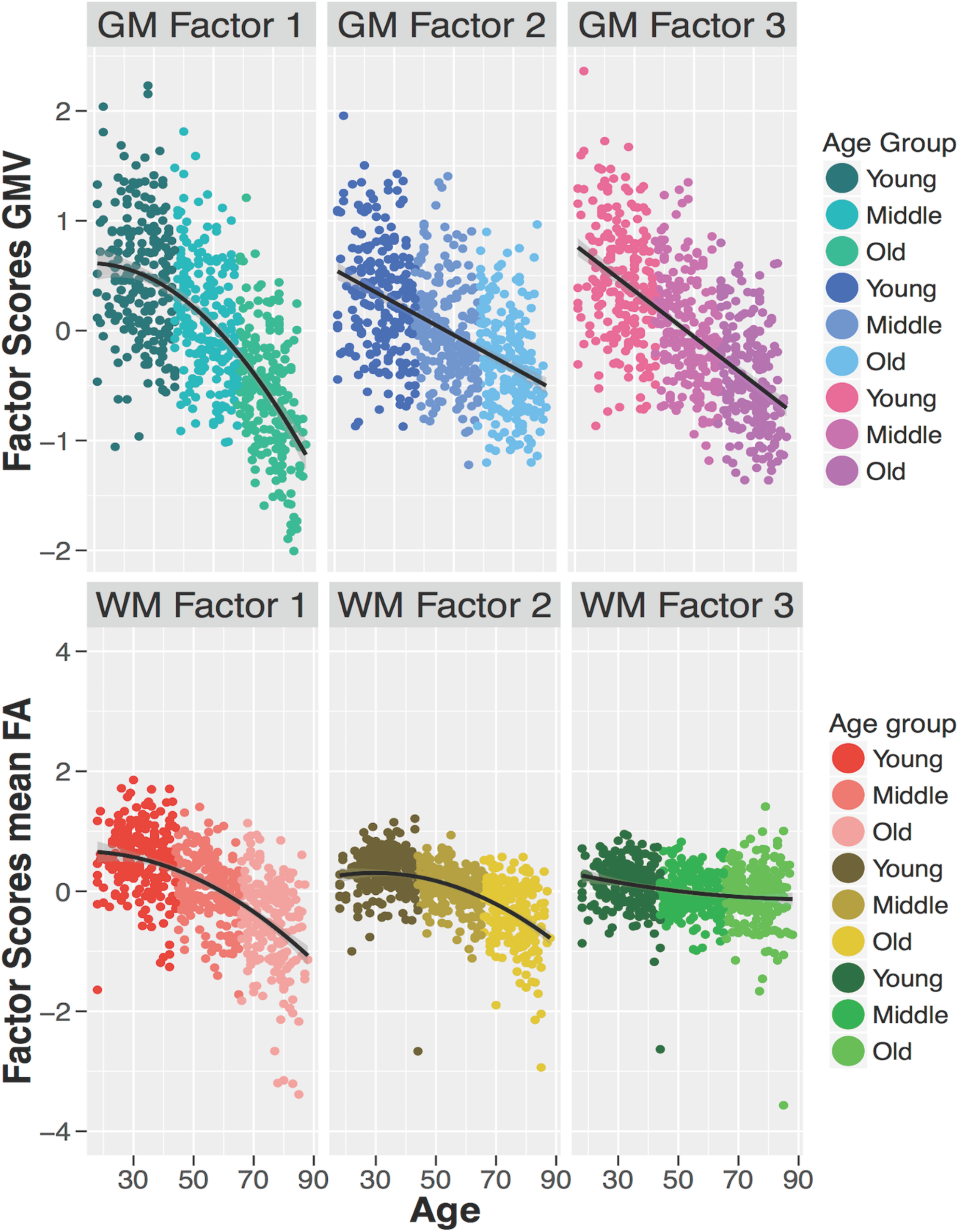
Age-related decline in grey matter (upper three plots) and white matter (lower plots) factor scores, according to the age groups (Young, Middle & Old) with best functional form shown (linear or non-linear). William’s test for dependent correlations showed that the effects of age were significantly different across the grey matter ROIs, t_cor1_2_ (651) = -9.46, *p* < .001; t_cor1_3_ (651) = -2.14, *p* = .033; t_cor2_3_ (651) = 12.79*,p* < .001, and across white matter ROIs, t_cor1_2_ (646) = -12.07, *p* < .001, t_cor1_3_ (646) = -12.07, *p* < .001, t_cor2_3_ (646) = – 8.28, *p* < .001.

First, we tested for age (de)differentiation using a multigroup Confirmatory Factor Analysis (MGCFA), where the population is divided into three age groups with equal sample sizes: young, middle & old (Table 1). This group-level comparison tests for age-related differences in specific parameters, while constraining the rest of the model (Δdf=6, as either three or nine factor covariances were estimated). Although constraints will generally lead to poorer model fit overall, we were interested in the specific comparison between the two nested models that represent age (de)differentiation versus no differentiation. Fitting these two models, we found that for the grey matter factors, a model where factor covariances were estimated freely across age groups showed better fit: Δ*χ^2^* (6) = 19.591, *p* = .003 (Table 2). Akaike weights showed that the freely-estimated model (with age-varying factor covariance) was 696 times more likely to be the better model given the data (Wagenmakers & Farrell, 2004). Using the same procedure for white matter, we found that the model with the freely estimated covariance also showed better fit: Δ*χ^2^* (6) = 25.430, *p* = .001, Akaike weights= 6297 in favor of the freely-estimated model.

**Table 2.**
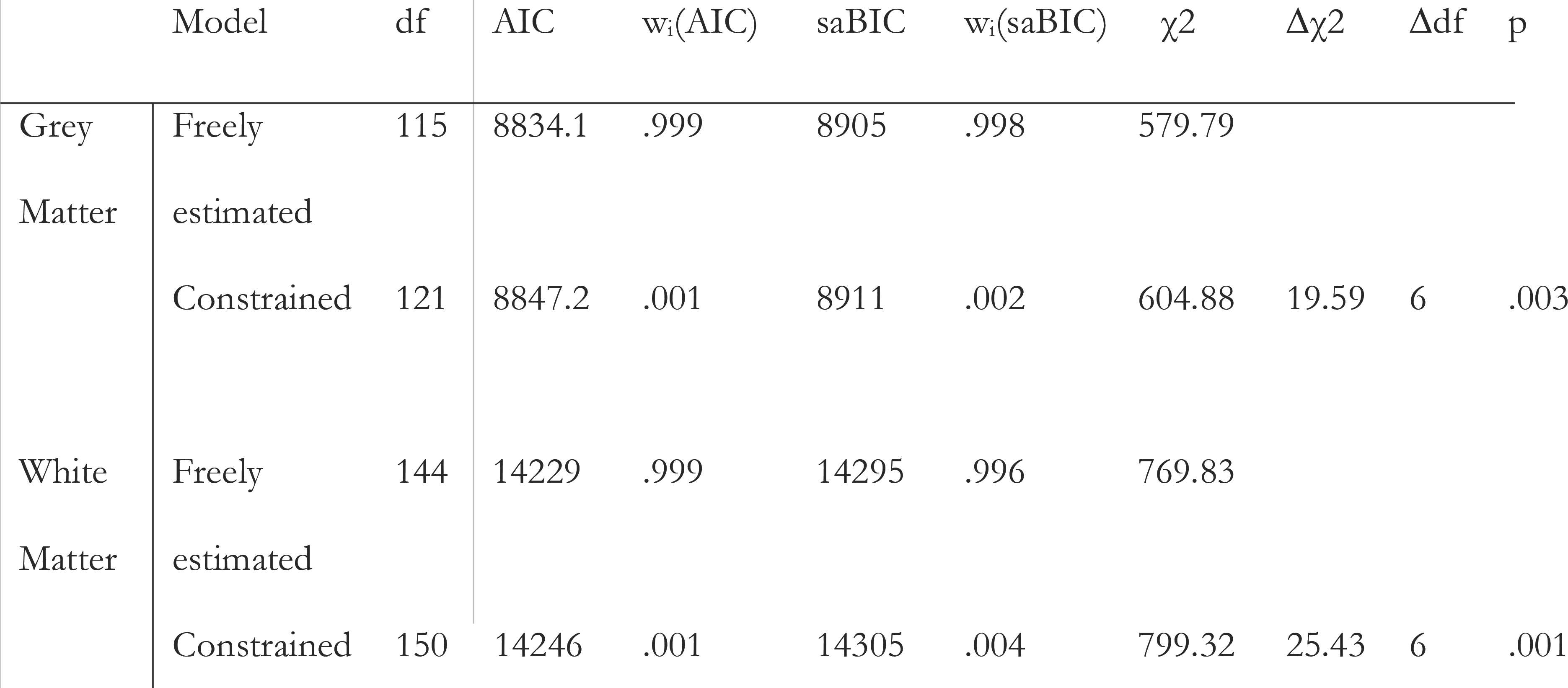
Model fit indices within white and grey matter, where the model with freely estimated covariance structure was preferred for both.

| | Model | df | AIC | $w_i(\text{AIC})$ | saBIC | $w_i(\text{saBIC})$ | $\chi^2$ | $\Delta\chi^2$ | $\Delta df$ | p |
| --- | --- | --- | --- | --- | --- | --- | --- | --- | --- | --- |
| Grey Matter | Freely | 115 | 8834.1 | .999 | 8905 | .998 | 579.79 |  |  |  |
|  | estimated |  |  |  |  |  |  |  |  |  |
|  | Constrained | 121 | 8847.2 | .001 | 8911 | .002 | 604.88 | 19.59 | 6 | .003 |
| White Matter | Freely | 144 | 14229 | .999 | 14295 | .996 | 769.83 |  |  |  |
|  | estimated |  |  |  |  |  |  |  |  |  |
|  | Constrained | 150 | 14246 | .001 | 14305 | .004 | 799.32 | 25.43 | 6 | .001 |

Next, we visualized the changing covariance within grey and white matter to assess evidence for age differentiation, de-differentiation, or some other pattern. The upper three plots in Figure 4 illustrate the difference in standardized covariance between each pair (GM1-GM2, GM1-GM3 and GM2-GM3) of grey matter factors. The strongest pattern is that factor GM1 displays considerable age differentiation: GM1 becomes more dissimilar to the two other grey matter factors with increasing age. For the white matter factors, the dominant pattern in the lower three plots of Figure 4 is the differentiation between factors WM1 and WM3, while the standardized covariance between factors WM1 and WM2, and between factors WM2 and WM3, remains relatively stable.

**Figure 4.**
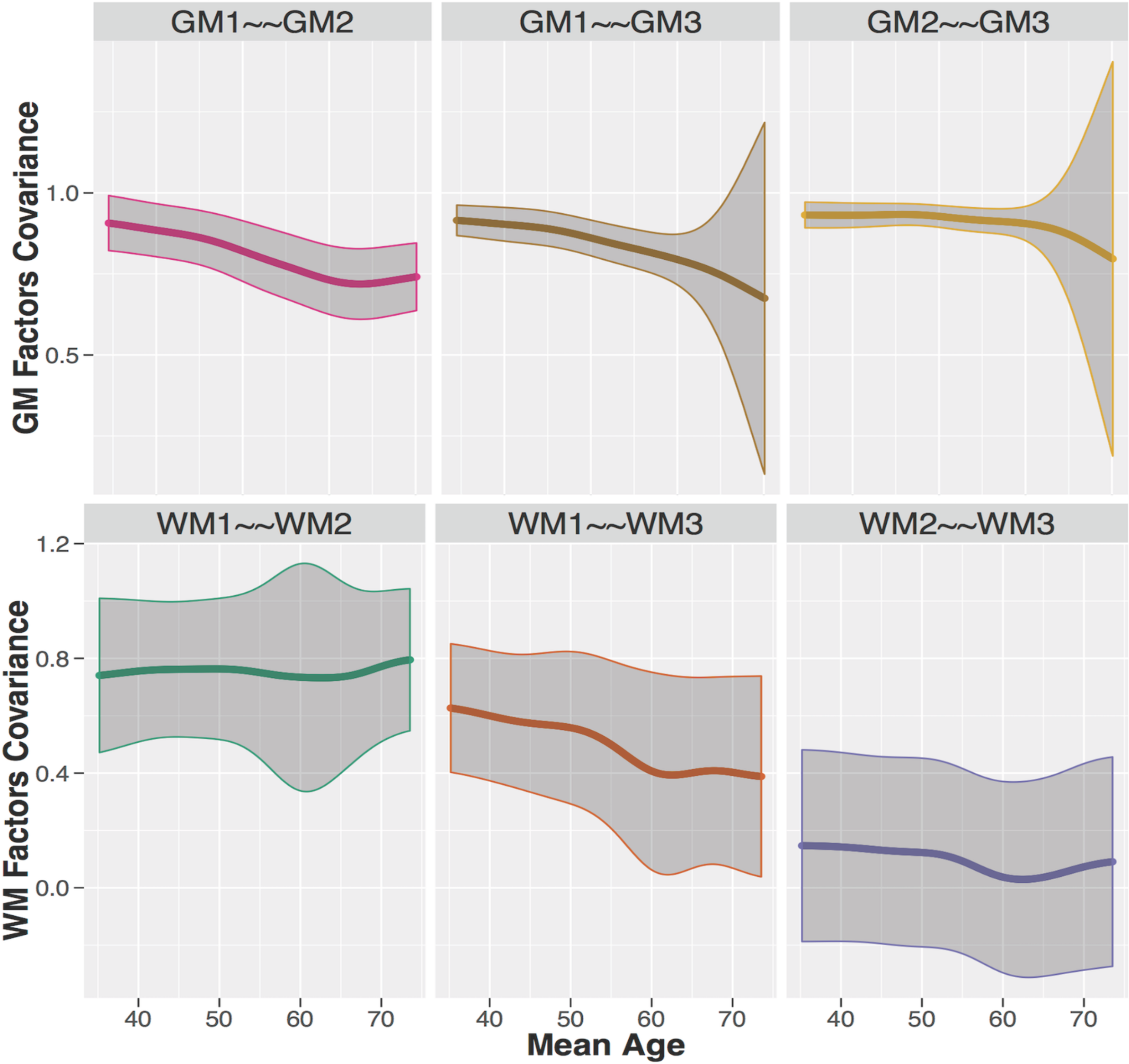
Differences in standardized covariance between the grey matter factors (top) and white matter factors (bottom) with age. 95% confidence intervals are displayed as the shaded area around die mean.

Finally, we validated the same question using the more exploratory technique of SEM trees with age as continuous covariate. For grey matter, the best split of the sample was given at the age of 50.5 years (*χ^2^* = 76.02, df=3), separating the participants into a young (N=285) and old (N=366) subgroup (left plot, figure 5). In line with the MGCFA, this analysis shows that the covariance between the grey matter factors decreases in old age. For white matter, we also find a single optimal split at a much older age of 66.5 years (χ^2^ = 36.07, df=3), separating participants in a younger (N=442) and older age group (N=204). The factor covariance between the white matter factors decreased in old age similar to grey matter (right plot, figure 5). Together, these three analytic strategies converge on the same conclusion: We observe age differentiation, or decreased covariance, among neural factors starting after middle age.

**Figure 5.**
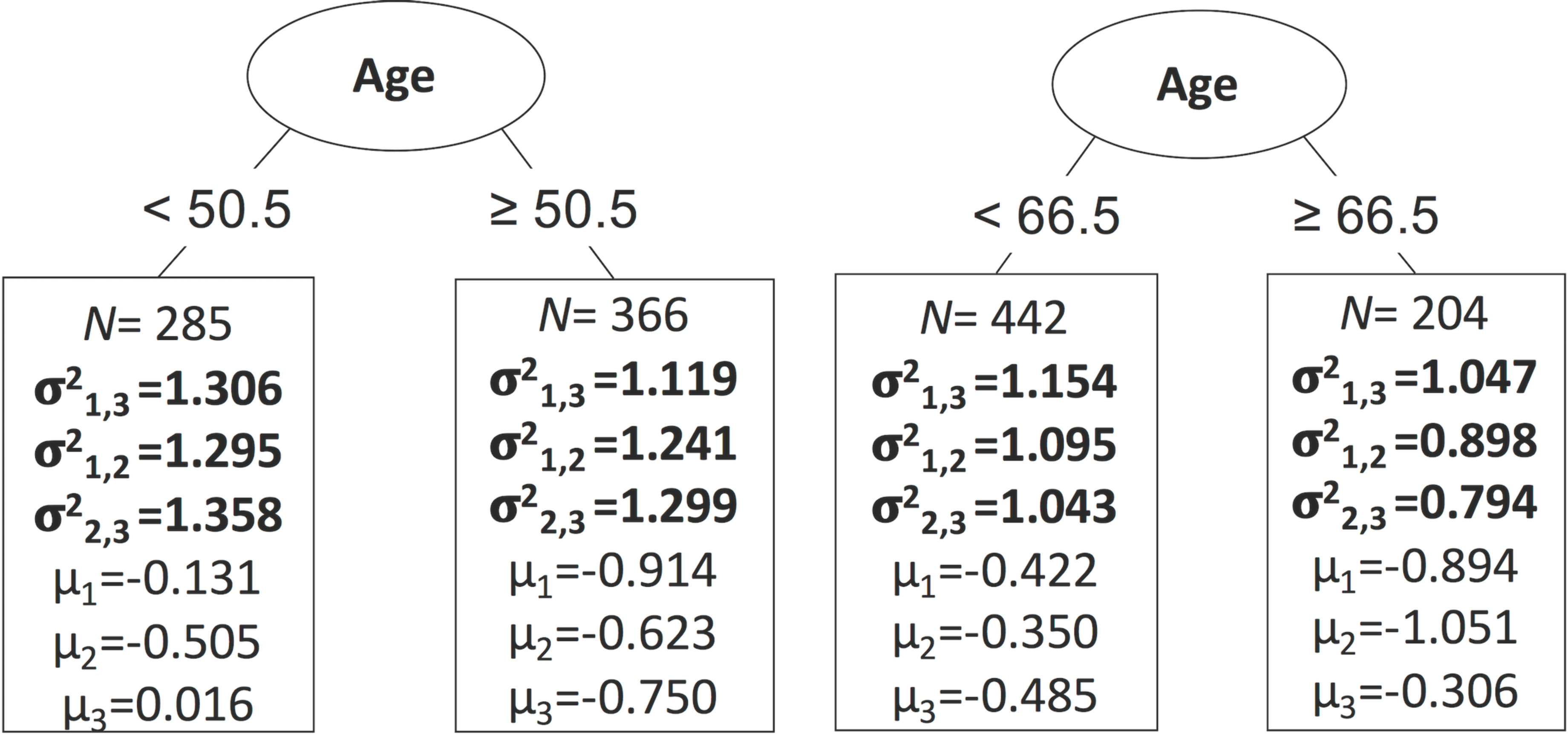
SEM Tree analysis with optimal splits for grey matter (left) at the age of 50.5 and for white matter (right) at the age of 66.5 years old. The standardized factor covariance (σ^2^) and factor means (μ) are depicted per subgroup, including the size of the group

A recent paper by Cox et al. (2016) employed a different analytic strategy than ours: Instead of focusing on factor covariance, they imposed a single factor model and examined factor loadings as they changed across the lifespan. To examine the robustness of our findings to such alternative approaches, we likewise imposed a single factor model across all brain regions, and tested whether factor loadings, rather than covariance, differ across three age groups (Young, Middle & Old). For grey matter, even though the one factor model did not fit well (χ^2^ (27) = 320.516, p < .001, RMSEA = .129 [.117 .141], CFI = .955, SRMR = .026), and a likelihood ratio test showed that it was a worse description of the data than the three-factor model (Δχ^2^ (8) = 227.66, p <.001), the model with freely-estimated factor loadings is again the better than the constrained model (Δχ^2^ (16) = 109.27, p <.001, Akaike weights= 6.13 * 1022), supporting differences in grey matter factor loadings across the adult lifespan. A visual inspection of the smoothed LOSEM shows that all factor loadings decline with age, again in line with age differentiation (Figure 6). Together, this represents strong evidence for age differentiation for grey matter factors; a pattern that does not depend on the precise analytical method.

**Figure 6.**
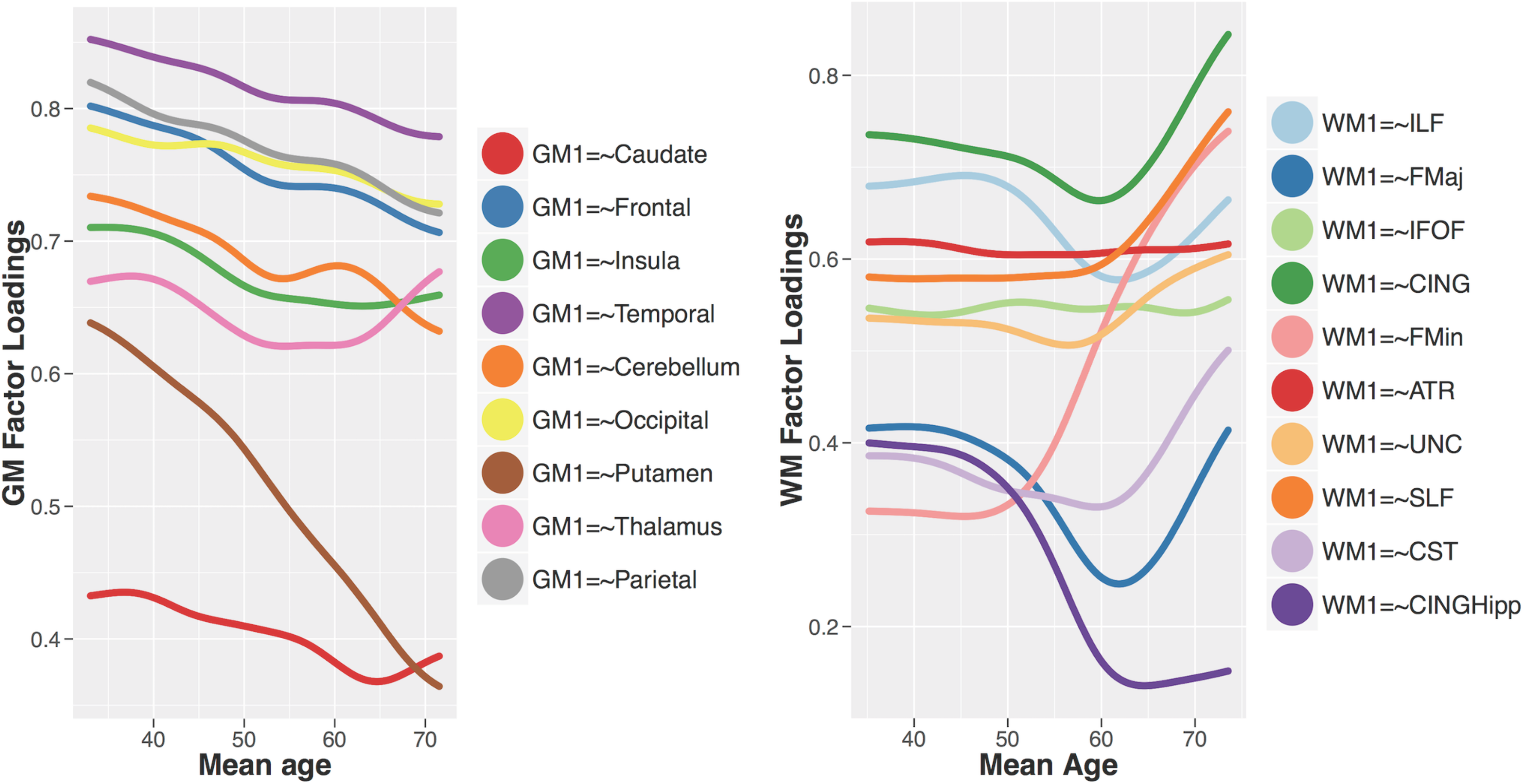
Standardized factor loadings in one factor model of grey matter (left) and white matter (right) across the lifespan

For the white matter, we again found that the one factor model for white matter did not fit well (χ^2^ (35) = 418.652, p < .001, RMSEA = .130 [.120 .140], CFI = .879, SRMR = .062), with the three-factor model showing better fit (Δχ^2^ (9) = 259.23, p <.001; Table 2). Nonetheless, within the single factor conceptualization, we again observe that the freely-estimated factor loadings were preferred over the constrained version (Akaike weights = 7.90 * 1028). The LOSEM plot in Figure 6 shows a complex pattern, with several factor loadings increasing (e.g. Forceps Minor and Superior Longitudinal Fasciculus), while others remain stable (e.g. Inferior Fronto-Occipital Fasciculus, Anterior Thalamic Radiations) or decline (e.g. Hippocampal Cingulum). The subset of increasing factor loadings is partly in line with Cox et al. (2016), who suggested age dedifferentiation of white matter tracts as the role of the general factor increases with age. However, the poor fit of the one-factor model, and the fact that factor loadings in our sample show both evidence for age differentiation as well as de-differentiation, suggest a cautious interpretation is warranted, with further, ideally longitudinal, investigation being crucial to understand the complex age-related differences in white matter covariance.

Finally, we implemented MGCFA on the combination of white and grey matter, with the same measurement models imposed, to see if the covariance between white and grey matter factors changes across the lifespan. We did not find evidence for age-related difference in the covariance between WM and GM: The more parsimonious constrained model of the covariance structure was more likely (Δχ^2^ (18) = 24.10, p = 0.152). These tests establish that the covariance within neural factors for both grey and white matter is different across the three age groups, but the covariance between the two neural measures does not differ across age groups.

### 3.2 Cognitive factors

We next examined age-related differences in covariance across cognitive factors. We defined three latent factors for the measurement model (see Figure 7A), based on a priori defined cognitive domains: (1) language, modelled by two Spot-the-Words tasks as a first-order factor and single proverb comprehension task, (2) fluid intelligence, fit to the four scores on Cattell subtests and (3) memory, fit to immediate recall, delayed recall and delayed recognition scores. The three-cognitive-factor model, shown in Figure 7A, fit the data well: *χ^2^* (31) = 59.030, p = .002, RMSEA = .036 [.022 .049], CFI = .988, SRMR = .030. The three-factor model fit considerably better than a one-factor solution (Δχ^2^ (4) = 336.43, p < 0.001; Akaike weights: 3.31 * 10273). Figure 7B shows the lifespan differences in the three cognitive factor scores.

**Figure 7.**
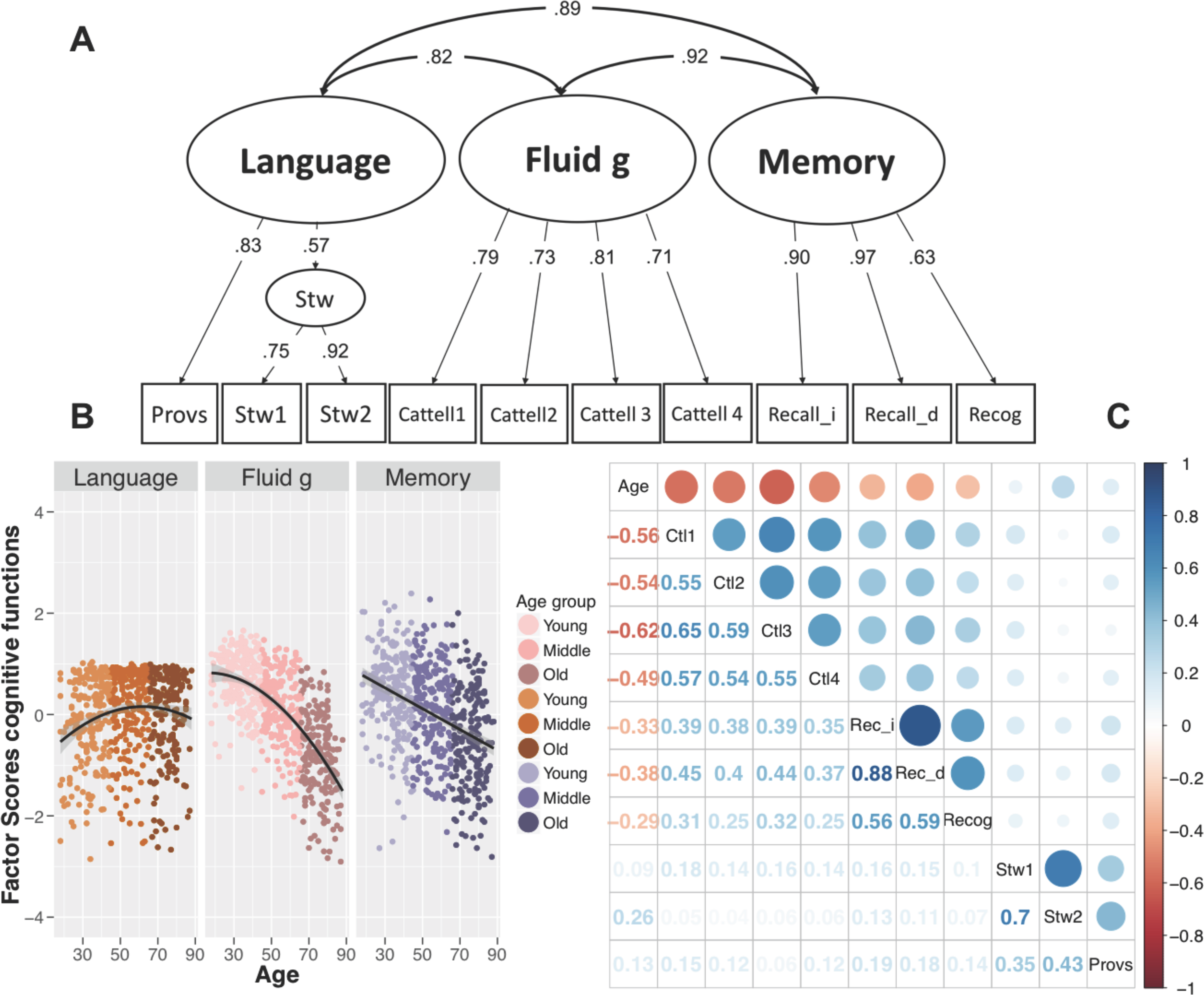
(A) Confirmatory factor model for cognitive processing based on Proverb Comprehension (Provs), two spot-the-word tasks (Stw1 and Stw2), four Catell subtests (Catell 1-4) relating fluid intelligence (Fluid g), immediate and delayed recall (Recall_i, Recall_d) and delayed recognition (Recog). All paths are fully standardized. (B) Age-related difference according to the age groups of the three cognitive factors: Language, Fluid intelligence and Memory, with best functional form shown (linear or nonlinear). William’s test for dependent correlations showed that the effects of age were significantly different between language and fluid g: t(707) = 24.21, *p* < .001; between fluid g and memory *t* (707) = -11.07, *p* < .001; and between language and memory: : t(707) = 12.9, *p* < .001. (C) correlation matrices between all cognitive tasks and age.

We looked for evidence for age differentiation among the cognitive factors across the three age groups with MGCFA, and found that the constrained covariance model was more likely: Δχ^2^ (6) = 4.984, p = 0.546, in line with an absence of either age-related cognitive differentiation or dedifferentiation. When we examined the same question using SEM Trees, we did not observe a significant split in covariance structure with age. The lack of evidence for (de)differentiation in both methods suggests a relative static covariance structure of cognitive abilities across the lifespan, contrary to studies of e.g. De Frias et al. (2007), but in line with Deary et al. (1996), Juan-Espinosa et al. (2002) and Tucker-Drob (2009).

### 3.3 Neurocognitive age differentiation

Finally, having examined brain and cognitive differentiation separately, we investigated their interaction to explore differences in brain-cognition covariance across the lifespan. To do so, we imposed the same measurement models as used above, first for grey matter and cognition, then for white matter and cognition. Our goal was to see if there is evidence for neurocognitive age (de)differentiation, indicated by differing covariance between brain structure and cognitive function across the lifespan.

With MGCFA we did not find evidence for neurocognitive age (de) differentiation in the covariance of grey matter with cognition: The more parsimonious constrained model of the covariance structure was preferred (Δχ^2^ (18) = 21.53, p = 0.253), suggesting a relative stable relationship between the grey matter and cognitive factors across the lifespan. Similarly, the SEM trees did not show a significant split in the factor covariance with age.

For white matter however, the multigroup analysis suggested that the freely estimated covariance structure was preferred: Δχ^2^ (18) = 37.27, p = .005, showing age-related differences in the relationship between white matter and cognitive factors. In the SEM tree analysis, we found an optimal split at the age of 56.5 years (χ2 = 60.15, df=9). Notably, all factor covariance in old age (N=335) decreased in comparison to the young age subgroup (N=372; Figure 8A).

To examine the source and trend of this neurocognitive age (de)differentiation, we plotted smoothed LOSEM age-weighted measurement models of the nine covariances between the three cognitive and three white matter factors (Figure 8B). Visual inspection suggested that this age-related difference in the relationship between cognition and white matter was driven most strongly by a specific pathway, namely the covariance between WM3 and memory. This visual inspection was confirmed by a post-hoc test, where a model with a freely-estimated covariance between WM3 and memory was strongly preferred over the constrained model: Δχ^2^ (2) = 27.34, p = <.001. The covariance between this white matter factor and memory performance declined steadily, especially in old age, suggesting a form of neurocognitive age differentiation. Further post-hoc comparisons for the other factors were not significant. It is noteworthy that the third white-matter factor was the (only) factor characterized by the ventral cingulum (CINGHipp), the part of the cingulum that is directly interconnected with the hippocampal formation (Hua et al. 2008, Fig. 2). This suggests a decoupling of memory performance from the white matter networks associated with the hippocampus; an intriguing pattern that we return to below.

**Figure 8.**
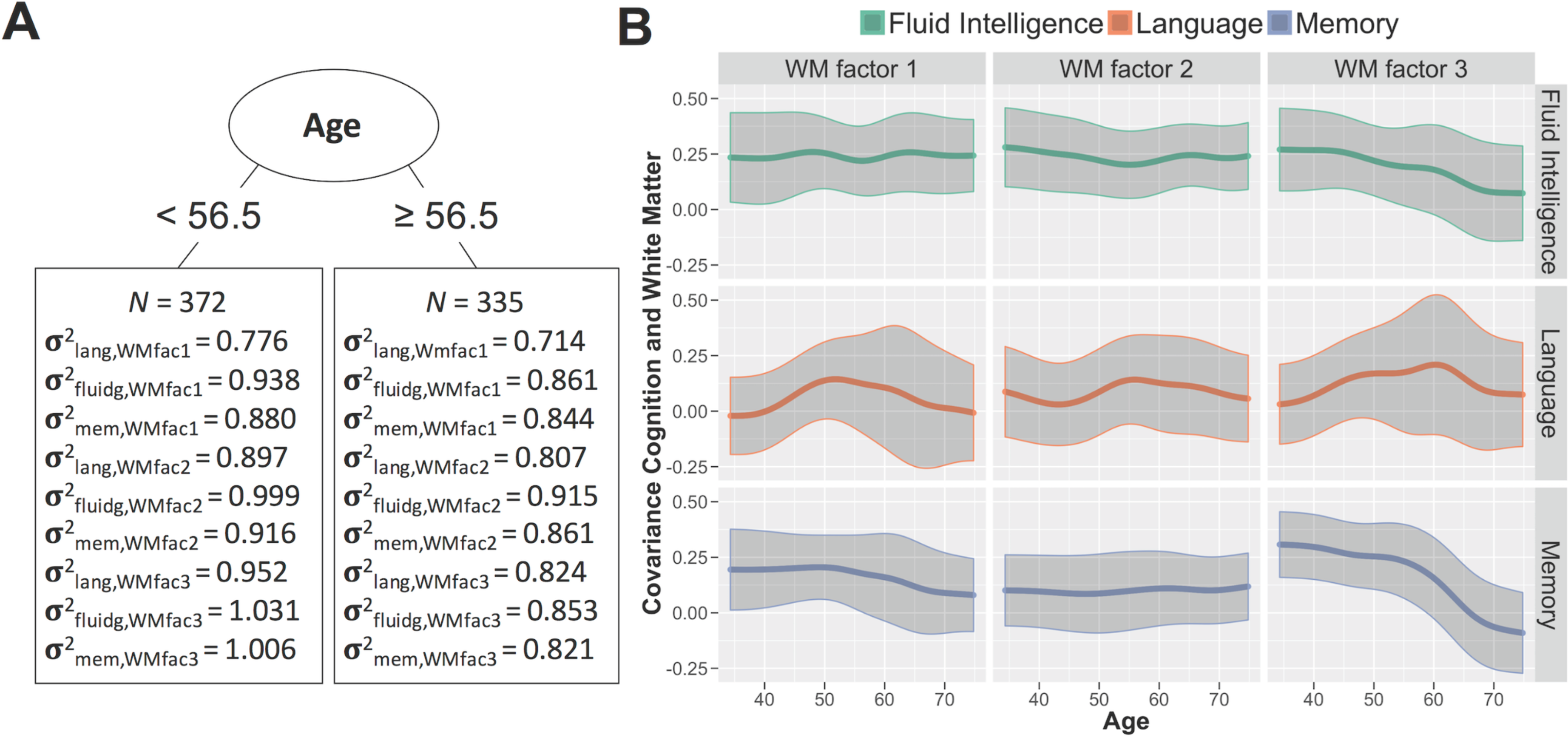
A) The optimal age split based on the factor covariance between white matter and the cognitive factors language (lang), fluid intelligence (fluidg) & memory (mem) using SEM Trees. B) Differences in the inter correlations between the cognitive and white matter factors across the lifespan according to LOSEM. The bottom right shows the one pathway that displays evidence for neurocognitive age differentiation.

## 4. Discussion

In this study, we examined the notion of age (de)differentiation within and between cognitive and neural factors across the adult lifespan. We found evidence for age differentiation within both GM and WM, such that the covariance between (a subset of) GM factors and the covariance between (a subset of) WM factors is lower in older adults. In contrast, the cognitive factors displayed a stable covariance structure, providing no evidence for (de)differentiation. Finally, we observed a specific pattern of age differentiation between WM and cognition, driven almost exclusively by a decoupling between a WM factor highly loading on the hippocampal cingulum, and the cognitive factor associated with memory.

For GM, exploratory factor analysis revealed that a three-factor model was preferred. The main effect of age was to reduce the covariance between the first factor (which loaded most on caudate and insula) and the other two factors. This neural differentiation was also observed when imposing a single-factor model, with factor loadings decreasing across the lifespan. Note that the precise number and nature of factors is likely to depend on the dimensionality of the data. Here we chose a mask characterized by a small number of ROIs (nine in total), in order to keep the GM model comparable in dimensionality to the white-matter tracts and cognitive variables. Moreover, a limited number of ROIs was necessary to achieve tractable SEM complexity given the sample size and subgroup analyses. Nonetheless, our ROIs had sufficient resolution to suggest that distinct networks of those regions differentiate in unique ways, resulting in structural networks that become more dissimilar across individuals in old age.

For WM, a three-factor model of the ten major WM tracts was also preferred. With this model, we again found evidence for differentiation, with the most noticeable effect being age-related reductions in the covariance between the first factor (which loaded most highly on the inferior fronto-occipital fasciculus and inferior longitudinal fasciculus) and the third factor (which loaded most highly on the ventral cingulum and projection fibers of corticospinal tract). The results from fitting an alternative single-factor model (e.g. Cox et al., 2016) were less clear, with both deceases and increases in various factor loadings with age, with the increases suggesting some dedifferentiation. A promising future avenue to better understand this complex pattern of white matter covariance differences is to examine longitudinal changes in white matter covariance, although at present there are few such datasets available.

Several mechanisms might contribute to our findings of differentiation within GM and within WM. First, the differentiation may reflect declines in structural connectivity during healthy ageing (Spreng & Turner, 2013). For example, reductions in grey matter covariance may follow reductions in white matter covariance (e.g. myelination) that cause less efficient communication and coactivation between brain regions, over time leading to decreased structural similarity. This is consistent with the present lack of evidence for differentiation between GM and WM. Another possibility is that the differentiation reflects distinct subpopulations of people that diverge across the lifespan. For instance, if subsets of the older population suffer from medical conditions that differentially affect specific brain regions (e.g. higher blood pressure, Gianaros et al., 2006), this will also lead to a more complex covariance pattern for the older population. Note that it is also possible that systemic age-related effects lead to age-related increases in covariance, or dominance of a single factor (Cox et al., 2016), which may be disguised by the causes of differentiation described above. Future studies should combine longitudinal imaging approaches with repeated health data to test the plausibility of these explanations in explaining the patterns observed here.

In line with most previous findings (e.g. Deary et al., 1996; Juan-Espinosa, 2002; Tucker-Drob, 2009), we did not observe evidence for cognitive age differentiation or dedifferentiation, instead finding a stable covariance structure across the lifespan. More importantly, we examined, for the first time, age differentiation between neural and cognitive factors. Specifically, we observed decreased covariance between a WM factor associated with hippocampal connectivity and a factor associated with memory. This decreased dependency of memory performance on WM integrity may relate to recent analyses of functional connectivity in healthy ageing. For instance, Salami, Pudas and Nyberg (2014) observed greater connectivity within a hippocampal network during rest in older relative to younger people, but decreased connectivity between the hippocampal network and other cortical networks during mnemonic tasks. Notably, this pattern of ‘aberrant hippocampal decoupling’ (p. 17654) was stronger in individuals with lower white matter integrity near the hippocampus, and was associated with poorer memory performance. Westlye et al. (2011) also found aberrant hippocampal functional connectivity associated with poorer performance, and suggested that failures of task-related hippocampal decoupling may elevate the risk of cognitive decline by increasing the metabolic burden on the hippocampus. In a longitudinal structural investigation, Gorbach et al. (2017) observed a robust brain-cognition change-change association between episodic memory decline and hippocampal atrophy in older adults (60-85 years), in line with brain maintenance. Future work integrating longitudinal investigations of the between-individual measurement models across time points in concert with within-subject change-change modelling will be able to reconcile these findings.

An alternative explanation of the decreased covariance between WM and memory observed here is the notion of cognitive reserve (Stern 2002, 2009; Whalley et al., 2004), which posits that the degree of brain pathology in certain individuals does not directly correspond to the manifestation of cognitive impairment. Certain lifespan exposures (e.g. high levels of education) are considered protective against cognitive decline. This implies that in older age, as the compensatory mechanisms of cognitive reserve become more prominent, memory performance should depend less on the brain structure, leading to the type of neurocognitive differentiation (i.e. decreased covariance) observed here. However, the precise consequences of cognitive reserve on covariance patterns likely depend on the idiosyncrasies of the sample under investigation. Moreover, it is unclear why we observe a mostly specific pattern of age-related differentiation (between WM and memory), rather than a more general neurocognitive differentiation.

A limitation of our study is that the sample is cross-sectional. The consequence is that, although we can examine age differentiation between individuals, we cannot generalize our findings to intraindividual changes over the lifespan (Salthouse, 2011). Acquiring longitudinal imaging and cognitive data would allow more detailed investigation of age-related changes in covariance among cognitive and neural factors. Moreover, the recruitment procedure in the Cam-CAN study included two age-correlated selection criteria that may bias the covariance population parameters: the exclusion of participants by general practitioners, and our exclusion of individuals with poor hearing and poor vision for reasons of procedural uniformity. Both hearing and vision are known to correlate with cognition, especially in old age (Baltes & Lindenberger, 1997), so that these procedures induce a positive selection bias of disproportionately healthy individuals in old age. Although age-correlated selection bias will inevitably be present in studies, the degree of bias can be reduced through alternative recruitment procedures such as general registry (e.g. de Frias et al., 2007) and/or using more liberal inclusion criteria such as in the Berlin Aging Study (Baltes & Lindenberger, 1997), where subgroups were blind or deaf or diagnosed with mild dementia. Furthermore, we focus on a relatively limited range of cognitive and neural variables, in order to enable SEMs with a tractable set of parameters. Possible solutions may be found in, for instance, regularized SEM (Jacobucci, Grimm, & McArdle, 2016) that allows measurement and structural models to be based on a larger set of neural and cognitive indicators. Alternatively, larger samples, possibly depending on integration across cohorts, would allow fitting of higher dimensional measurement models (with possibly an overall better fit) and simultaneously explore generalizability. A second limitation of our study concerns potential differences in data quality across the lifespan. For instance, if older adults move more, and the effects of this motion of the imaging data cannot be fully accommodated (Geerligs et al., 2017), this may induce a decrease in covariance simply due to less reliable measurement. However, age-related decreases in data quality would seem unlikely to fully explain our findings, given that the pattern of age differentiation was limited to some, but not all, neural factors: increased measurement error in older adults would be expected to produce more uniform decreases in covariance between all pairs of factors.

Our findings show how multigroup Confirmatory Factor Analysis and SEM Trees can be powerful techniques for investigating theories of neurocognitive ageing, such as age differentiation, allowing researchers to investigate mechanisms of healthy and pathological ageing in a flexible yet principled manner. Taken together, these techniques revealed a complex pattern of age-related differentiation in grey and white matter, but not in cognition, together with a specific differentiation in the relationship between white-matter tracts and memory. Future work on the long term, developmental patterns of covariance across the lifespan may help further elucidate the mechanisms underlying these observations.

## Acknowledgements

The Cambridge Centre for Ageing and Neuroscience (Cam-CAN) research was supported by the Biotechnology and Biological Sciences Research Council (grant number BB/H008217/1). This project has also received funding from the European Union’s Horizon 2020 research and innovation programme (grant agreement number 732592). The Cam-CAN team (see http://www.cam-can.com/) was crucial in recruiting participants, developing the protocol, conducting the testing and overseeing data management. R.N.A.H. was supported by MRC Programme Grant MC-A060-5PR10. RAK is supported by the Sir Henry Wellcome Trust (grant number 107392/Z/15/Z) and MRC Programme Grant MC-A060-5PR60. We are grateful to the Cam-CAN respondents and their primary care team in Cambridge for their participation in this study. We also thank colleagues at the MRC Cognition and Brain Sciences Unit MEG and MRI facilities for their assistance.

## Cam-CAN Research team

Lorraine K. Tyler^4^, Carol Brayne^4^, Ed Bullmore^4^, Andrew Calder^4^, Rhodri Cusack^4^, Tim Dalgleish^4^, Fiona Matthews^4^, William Marslen-Wilson^4^, James Rowe^4^, Meredith Shafto^4^; Research Associates: Karen Campbell^4^, Teresa Cheung^4^, Linda Geerligs^4^, Anna McCarrey^4^, Kamen Tsvetanov^4^, Nitin Williams^4^; Research Assistants: Lauren Bates^4^, Tina Emery^4^, Sharon Erzinc, lioglu^4^, Andrew Gadie^4^, Sofia Gerbase^4^, Stanimira Georgieva^4^, Claire Hanley^4^, Beth Parkin^4^, David Troy^4^; Research Interviewers: Jodie Allen^4^, Gillian Amery^4^, Liana Amunts^4^, Anne Barcroft^4^, Amanda Castle^4^, Cheryl Dias^4^, Jonathan Dowrick^4^, Melissa Fair^4^, Hayley Fisher^4^, Anna Goulding^4^, Adarsh Grewal^4^, Geoff Hale^4^, Andrew Hilton^4^, Frances Johnson^4^, Patricia Johnston^4^, Thea Kavanagh-Williamson^4^, Magdalena Kwasniewska^4^, Alison McMinn^4^, Kim Norman^4^, Jessica Penrose^4^, Fiona Roby^4^, Diane Rowland^4^, John Sargeant^4^, Maggie Squire^4^, Beth Stevens^4^, Aldabra Stoddart^4^, Cheryl Stone^4^, Tracy Thompson^4^, Ozlem Yazlik^4^; and Administrative Staff: Dan Barnes^4^, Marie Dixon^4^, Jaya Hillman^4^, Joanne Mitchell^4^, Laura Villis^4^

## Footnotes

4 Cambridge Centre for Ageing and Neuroscience (Cam-CAN), University of Cambridge and MRC Cognition and Brain Sciences Unit, Cambridge CB2 7EF, UK.

## References

Alexander-Bloch, A., Giedd, J. N., & Bullmore, E. (2013). Imaging structural covariance between human brain regions. Nature Reviews Neuroscience, 14, 322–336.

Alexander-Bloch, A., Raznahan, A., Bullmore, E., & Giedd, J. (2013). The convergence of maturational change and structural covariance in human cortical networks. The Journal of Neuroscience, 33, 2889–2899.

Arshad, M., Stanley, J. A., & Raz, N. (2016). Adult age differences in subcortical myelin content are consistent with protracted myelination and unrelated to diffusion tensor imaging indices. NeuroImage, 143, 26–39.

Baddeley, A., Emslie, H., & Nimmo-Smith, I. (1993). The Spot-the-Word test: A robust estimate of verbal intelligence based on lexical decision. British Journal of Clinical Psychology, 32, 55–65.

Baltes, P. B., & Lindenberger, U. (1997). Emergence of a powerful connection between sensory and cognitive functions across the adult life span: a new window to the study of cognitive aging? Psychology and aging, 12, 12.

Blum, D., & Holling, H. (2017). Spearman’s law of diminishing returns. A meta-analysis. Intelligence.

Brandmaier, A. M., Prindle, J. J., McArdle, J. J., & Lindenberger, U. (2016). Theory-guided exploration with structural equation model forests. Psychological methods, 21, 566.

Brandmaier, A. M., von Oertzen, T., McArdle, J. J., & Lindenberger, U. (2013). Structural equation model trees. Psychological methods, 18, 71.

Cabeza, R., Anderson, N. D., Locantore, J. K., & McIntosh, A. R. (2002). Aging gracefully: compensatory brain activity in high-performing older adults. Neuroimage, 17, 1394–1402.

Cattell, R. B. (1971). Abilities: Their structure, growth, and action

Cox, S. R., Ritchie, S. J., Tucker-Drob, E. M., Liewald, D. C., Hagenaars, S. P., Davies, G.,… & Deary, I. J. (2016). Ageing and brain white matter structure in 3,513 UK Biobank participants. Nature communications, 7, 13629.

Damoiseaux, J. S., & Greicius, M. D. (2009). Greater than the sum of its parts: a review of studies combining structural connectivity and resting-state functional connectivity. Brain Structure and Function, 213, 525–533.

Deary, I. J., Egan, V., Gibson, G. J., Austin, E. J., Brand, C. R., & Kellaghan, T. (1996). Intelligence and the differentiation hypothesis. Intelligence, 23, 105–132.

Deary, I. J., & Pagliari, C. (1991). The strength of g at different levels of ability: Have Detterman and Daniel rediscovered Spearman’s “law of diminishing returns”? Intelligence, 15, 247–250.

de Frias, C. M., Lövdén, M., Lindenberger, U., & Nilsson, L. G. (2007). Revisiting the dedifferentiation hypothesis with longitudinal multi-cohort data. Intelligence, 35, 381–392.

Di, X., Gohel, S., Thielcke, A., Wehrl, H. F., Biswal, B. B., & Alzheimer’s Disease Neuroimaging Initiative. (2017). Do all roads lead to Rome? A comparison of brain networks derived from inter-subject volumetric and metabolic covariance and moment-to-moment hemodynamic correlations in old individuals. Brain Structure and Function, 1–13.

Garrett, H. E. (1946). A developmental theory of intelligence. American Psychologist, 1, 372.

Geerligs, L., Tsvetanov, K., Cam-CAN & Henson, R.N. (2017). Challenges in measuring individual differences in functional connectivity using fMRI: the case of healthy aging. Human Brain Mapping, 38, 4125–4156.

Ghisletta, P., & Lindenberger, U. (2003). Age-based structural dynamics between perceptual speed and knowledge in the Berlin Aging Study: direct evidence for ability dedifferentiation in old age. Psychology and aging, 18, 696.

Gianaros, P. J., Greer, P. J., Ryan, C. M., & Jennings, J. R. (2006). Higher blood pressure predicts lower regional grey matter volume: Consequences on short-term information processing. Neuroimage, 31, 754–765.

Gorbach, T., Pudas, S., Lundquist, A., Orädd, G., Josefsson, M., Salami, A.,… & Nyberg, L. (2017). Longitudinal association between hippocampus atrophy and episodic-memory decline. Neurobiology of aging, 51, 167–176.

Greenwood, P. M. (2007). Functional plasticity in cognitive aging: review and hypothesis. Neuropsychology, 21(6), 657.

Hildebrandt, A., Wilhelm, O., & Robitzsch, A. (2009). Complementary and competing factor analytic approaches for the investigation of measurement invariance. Review of Psychology, 16, 87–102.

Honey, C. J., Sporns, O., Cammoun, L., Gigandet, X., Thiran, J. P., Meuli, R., & Hagmann, P. (2009). Predicting human resting-state functional connectivity from structural connectivity. Proceedings of the National Academy of Sciences, 106, 2035–2040.

Hua, K., Zhang, J., Wakana, S., Jiang, H., Li, X., Reich, D. S.,… & Mori, S. (2008). Tract probability maps in stereotaxic spaces: analyses of white matter anatomy and tract-specific quantification. Neuroimage, 39, 336–347.

Hülür, G., Wilhelm, O., & Robitzsch, A. (2011). Intelligence differentiation in early childhood. Journal of Individual Differences.

Jacobucci, R., Grimm, K. J., & McArdle, J. J. (2016). Regularized structural equation modeling. Structural equation modeling: a multidisciplinary journal, 23, 555–566.

Jones, D. K., & Cercignani, M. (2010). Twenty-five pitfalls in the analysis of diffusion MRI data. NMR in Biomedicine, 23, 803–820.

Jones, D. K., Knösche, T. R., & Turner, R. (2013). White matter integrity, fiber count, and other fallacies: the do’s and don’ts of diffusion MRI. Neuroimage, 73, 239–254.

Juan-Espinosa, M., Garcia, L. F., Escorial, S., Rebollo, I., Colom, R., & Abad, F. J. (2002). Age dedifferentiation hypothesis: Evidence from the WAIS III. Intelligence, 30, 395–408.

Kievit, R. A., Davis, S. W., Griffiths, J., Correia, M. M., & Henson, R. N. (2016). A watershed model of individual differences in fluid intelligence. Neuropsychologia, 91, 186–198.

Kochunov, P., Williamson, D. E., Lancaster, J., Fox, P., Cornell, J., Blangero, J., & Glahn, D. C. (2012). Fractional anisotropy of water diffusion in cerebral white matter across the lifespan. Neurobiology of aging, 33, 9–20.

Li, S. C., Lindenberger, U., Hommel, B., Aschersleben, G., Prinz, W., & Baltes, P. B. (2004). Transformations in the couplings among intellectual abilities and constituent cognitive processes across the life span. Psychological Science, 15, 155–163.

Lindenberger, U., & Mayr, U. (2014). Cognitive aging: is there a dark side to environmental support? Trends in Cognitive Sciences, 18, 7–15.

Mazziotta, J., Toga, A., Evans, A., Fox, P., Lancaster, J., Zilles, K.,… & Holmes, C. (2001). A probabilistic atlas and reference system for the human brain: International Consortium for Brain Mapping (ICBM). Philosophical Transactions of the Royal Society of London B: Biological Sciences, 356, 1293–1322.

Mechelli, A., Friston, K. J., Frackowiak, R. S., & Price, C. J. (2005). Structural covariance in the human cortex. The Journal of neuroscience, 25, 8303–8310.

Molenaar, D., Dolan, C. V., Wicherts, J. M., & van der Maas, H. L. (2010). Modeling differentiation of cognitive abilities within the higher-order factor model using moderated factor analysis. Intelligence, 38, 611–624.

Molenaar, D., Kő, N., Rózsa, S., & Mészáros, A. (2017). Differentiation of cognitive abilities in the WAIS-IV at the item level. Intelligence, 65, 48–59.

Nyberg, L., Lövdén, M., Riklund, K., Lindenberger, U., & Bäckman, L. (2012). Memory aging and brain maintenance. Trends in cognitive sciences, 16, 292–305.

Park, D. C., Polk, T. A., Park, R., Minear, M., Savage, A., & Smith, M. R. (2004). Aging reduces neural specialization in ventral visual cortex. Proceedings of the National Academy of Sciences of the United States of America, 101, 13091–13095.

Park, D. C., & Reuter-Lorenz, P. (2009). The adaptive brain: aging and neurocognitive scaffolding. Annual review of psychology, 60, 173.

R Core Team (2016). R: A language and environment for statistical computing. R Foundation for Statistical Computing, Vienna, Austria. URL: https://www.R-project.org/.

Rosseel, Y. (2012). lavaan: An R package for structural equation modeling. Journal of Statistical Software, 48, 1–36.

Salami, A., Pudas, S., & Nyberg, L. (2014). Elevated hippocampal resting-state connectivity underlies deficient neurocognitive function in aging. Proceedings of the National Academy of Sciences, 111, 17654–17659.

Salthouse, T. A. (2011). Neuroanatomical substrates of age-related cognitive decline. Psychological bulletin, 137, 753.

Schermelleh-Engel, K., Moosbrugger, H., & Müller, H. (2003). Evaluating the fit of structural equation models: Tests of significance and descriptive goodness-of-fit measures. Methods of psychological research online, 8, 23–74.

Seeley, W. W., Crawford, R. K., Zhou, J., Miller, B. L., & Greicius, M. D. (2009). Neurodegenerative diseases target large-scale human brain networks. Neuron, 62, 42–52.

Shafto, M. A., Tyler, L. K., Dixon, M., Taylor, J. R., Rowe, J. B., Cusack, R.,… & Henson, R. N. (2014). The Cambridge Centre for Ageing and Neuroscience (Cam-CAN) study protocol: a cross-sectional, lifespan, multidisciplinary examination of healthy cognitive ageing. BMC, neurology, 14, 1

Spearman, C. (1927). The abilities of man.

Spreng, R. N., & Turner, G. R. (2013). Structural covariance of the default network in healthy and pathological aging. The Journal of Neuroscience, 33, 15226–15234.

Stern, Y. (2002). What is cognitive reserve? Theory and research application of the reserve concept. Journal of the International Neuropsychological Society, 8, 448–460.

Stern, Y. (2009). Cognitive reserve. Neuropsychologia, 47, 2015–2028.

Taylor, J. R., Williams, N., Cusack, R., Auer, T., Shafto, M. A., Dixon, M.,… & Henson, R. N. (2017). The Cambridge Centre for Ageing and Neuroscience (Cam-CAN) data repository: structural and functional MRI, MEG, and cognitive data from a cross-sectional adult lifespan sample. Neuroimage, 144, 262–269.

Tsang, A., Lebel, C. A., Bray, S. L., Goodyear, B. G., Hafeez, M., Sotero, R. C.,… & Frayne, R. (2017). White matter structural connectivity is not correlated to cortical resting-state functional connectivity over the healthy adult lifespan. Frontiers in Aging Neuroscience, 9.

Tucker-Drob, E. M. (2009). Differentiation of cognitive abilities across the life span. Developmental psychology, 45, 1097.

Wagenmakers, E. J., & Farrell, S. (2004). AIC model selection using Akaike weights. Psychonomic bulletin & review, 11, 192–196.

Wandell, B. A. (2016). Clarifying human white matter. Annual review of neuroscience, 39, 103–128.

Wechsler, D. (1997). WAIS-III: Wechsler adult intelligence scale. Psychological Corporation.

Westlye, E. T., Lundervold, A., Rootwelt, H., Lundervold, A. J., & Westlye, L. T. (2011). Increased hippocampal default mode synchronization during rest in middle-aged and elderly APOE ε4 carriers: relationships with memory performance. Journal of Neuroscience, 31, 7775–7783.

Whalley, L. J., Deary, I. J., Appleton, C. L., & Starr, J. M. (2004). Cognitive reserve and the neurobiology of cognitive aging. Ageing research reviews, 3, 369–382.

Wolf, E. J., Harrington, K. M., Clark, S. L., & Miller, M. W. (2013). Sample size requirements for structural equation models an evaluation of power, bias, and solution propriety. Educational and Psychological Measurement, 73, 913–934.

Zelinski, E. M., & Lewis, K. L. (2003). Adult age differences in multiple cognitive functions: differentiation, dedifferentiation, or process-specific change? Psychology and aging, 18, 727.

Zielinski, B. A., Gennatas, E. D., Zhou, J., & Seeley, W. W. (2010). Network-level structural covariance in the developing brain. Proceedings of the National Academy of Sciences, 107, 18191–18196.

